# Comment to: “Topology of molecular deformations induces triphasic catch bonding in selectin–ligand bonds”

**DOI:** 10.1101/2024.08.21.608529

**Authors:** Wolfgang Quapp, Josep Maria Bofill

## Abstract

We contradict diverse mathematical claims of a paper by Casey O. Barkan and Robijn F. Bruinsma in PNAS 2024, 121, No. 6, e2315866121, former BioRxiv preprint from Sept.12,2023. It deals with the physical mechanisms of protein-ligand catch bonding for the family of selectin proteins. Selectins exhibit slip, catch–slip, and slip–catch–slip bonding.

---

We congratulate Barkan and Bruinsma (BB) for the fitting of some quite diverse models of potential energy surfaces (PES) for different selectins (1), see the left panel of Figure 1. The class of proteins show the strange behavior of catch bonds. This remark concerns mathematical errors. BB use parts of the theory of Newton trajectories (NT) (2, 3) as a tool to rationalize the biochemical phenomena of slip- and catch-bonds. They study with (4) the mechanochemical potential

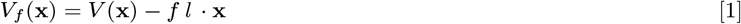

where *V* (.) is the PES, or the free energy surface, of a molecule, *l* is the direction of an external force vector acting on the molecule, and *f* is the magnitude of the force. Note that ansatz [1] is the simplest possible, with a linear external force. The stationary points of the PES move under the action of the force. For the movement of any critical point, x_c_, BB develop a differential equation, their Eq.[3], by

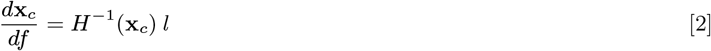

where *H*^−1^ is the inverse of the Hessian matrix of the original PES. We wonder about this differential equation. The determinant of *H* is positive in minima and negative in saddle points of index one (SP_1_) of the PES. So there is always a point on the way from a minimum to an SP_1_ where the determinant of *H* is zero and where Eq.[2] becomes singular. This problem was solved long ago by Branin (5). The better equation is given in reference 30 of BB citing one of our articles (2)

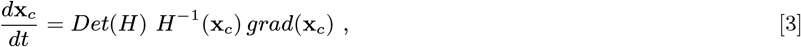

**Fig. 1.**
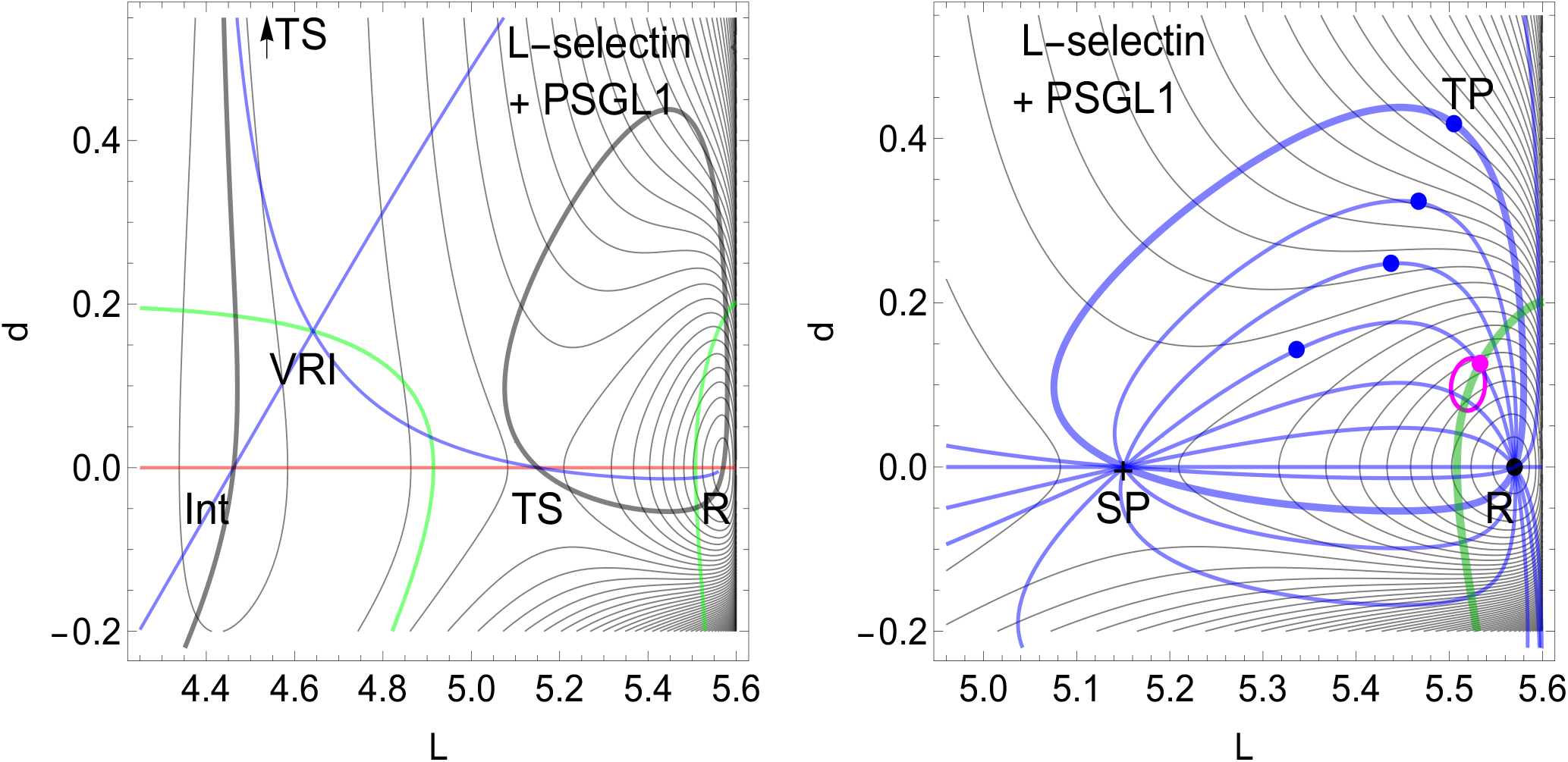
Left: Proposed PES of L-selectin (1). Coordinates are x=(L, d) in nm. Thin curves are level lines, the red line is the NT to direction (−1,0) – a clear slip bond direction. The thick black curve is the NT to direction (1,1), a direction with catch-slip behavior (1). The blue curve is the singular NT through the valley-ridge inflection (VRI) point forming the separatrix of the problem, and green curves are Det(H)=0 lines. Right: Part with different NTs (blue) and a trajectory to Eq.[2] (magenta). The upper four NTs have turning points (TP), blue dots. The magenta dot may indicate the crossing of the NT with the first TP with the green line. This can be an indicator for catch bonding. Note the magenta squiggle: it is the trajectory to Eq.[2] with start at the magenta dot. It is not an NT.

So the right one desingularized equation [3] is made worse into an equation with a singularity [2]. The solution curves of the Branin equation [3] are named Newton trajectories, since more than a third of a century (2, 3, 6). Every NT fulfills this Eq.[3].

## Methods and results

Our method is mathematical reasoning on the equations of the paper by BB (1), and the comparison with known properties of NTs (2, 3, 6–9). We obtain contradictory results to the claims of reference (1), see the right panel of Figure 1 below where the magenta squiggle line is a trajectory to Eq.[2]. It is not an NT. Detailed results are given in the following sections where especially in section ‘The field of NTs’ a new propose is discussed for a possible first emergence of catch bond behavior.

For Figure 1 we used Mathematica 13.3.1.0 for platform Linux x86 (64-bit) for calculations as well as for graphics.

### The equation [2]

The singularities of the equation are artificial, so to speak. And they do not generally generate a force-induces switch, like it is claimed in paper (1). Every solution of Eq. [3] to different directions *l* connecting a minimum with an SP_1_ crosses the curve of points where *Det*(*H*) = 0 applies (the green curves), so also right regular and ‘direct’ solutions. In the right panel of Figure 1 we give a family of NTs on the given PES. Every NT follows another direction of the gradient of the PES. The force in direction *l* with the special magnitude, *f*, to reach the *Det*(*H*) = 0 curve forces a coalescence of former minimum and former saddle SP_1_ for the effective PES. These points now form a shoulder on the effective PES [1]. This event is named bond breaking point (BBP) (10, 11). Each local point on the *Det*(*H*) = 0 curve determines one solution curve of Eq.[3], through its corresponding gradient direction there. The reason is that along each solution curve of Eq.[3] the gradient direction is fixed because it is equal to a fixed *l*. It means that one cannot start on different points on the *Det*(*H*) = 0 curve with the same direction *l*.

### The field of NTs

A remark to the ‘flow’ image in Figure 1D of the commented paper (1). This picture is misleading for the imagination of NTs. Every NT starts in a stationary point of the original PES, *V* (*x*), and a family of NTs leads to the next stationary point with an index difference of one (12). Each NT follows exactly one direction, *l*. Every NT of the family of different NTs in Figure 1, right panel, follows one gradient direction of the **gradient field** of the PES.

The arrow field like in Figure 1D of paper (1) offers a general meaning of the vectors of Eq. [2]. This is not correct. No, there is always a one-to-one relationship of different external directions, *l*, and different NTs. The field which the NTs follow is the field of constant gradient directions of the original PES.

An interesting property for catch bond behavior may be the emergence of turning points of corresponding NTs. We have indicated four TPs in Figure 1, right panel, by blue dots. May be one can use the first emergence of a TP for a hint to a possible switch in the slip- to catch bond character of NTs. For this we have indicated the magenta dot in Figure 1. NTs without a TP on their energy profile are clearly slip bonds (3), however, a TP is not sufficient for catch bonding, see an extreme counterexample in (2). We conclude that the *l*-switch points of BB in (1) are not well justified.

### The ‘ellipticity’ of field [2]

The upper remark also touches on the claimed ‘ellipticity’ near a so called *l*-switching point (1). The word implicates a circular behavior of NTs which is not true, in the general situation of a PES. In the right panel of Figure 1 we have included a solution trajectory of differential equation [2] (in magenta color) with start in the magenta dot, and the given fixed *l*=(1, 1) of [2]. It demonstrates well the circular character of the field of [2], however, it has nothing to do with the solution of our problem: the movement of stationary points of the PES under an external force. The direction *l*=(1, 1) of (1) counts only for the fat NT in our Figure 1. Besides this fat NT this direction is not of interest. To draw a vector field like in Figure 1D of (1) is not false, but it is useless for our problem.

### Further problems

We have to identify a further weakness in Eq.[3] of (1), here Eq.[2], the use of the force *f* for a curve length parameter. The Branin equation [3] uses an ‘extra’ curve length parameter, *t*, because *f* has to have on the way from minimum to SP a maximal value in between, at the corresponding BBP, compare many illustrations in ref. (2). Thus *f* is not the curve length parameter of the NT. However, *f* decreases to zero again to reach the SP.

The separatix for the catch bond behavior of an NT is a singular NT through the VRI point: because it is also an NT, it has to go through the TS on the L-axis and through the minimum, *R*, compare Figure 1. It is false in Figures of paper (1) where the separatrix misses these stationary points. BB also claim an equal topology of all selectin models. However, for E-selectin is missing the VRI point and the separatrix.

A remark to the *n* points of (1), the orange dots with number 2: they are the known and often discussed valley-ridge inflection points (VRI) (13) of the PES. They are characterized by a special kind of NTs, so called singular NTs which bifurcate at the VRI point. The singular NT through the bifurcation then has four branches. From minimum one to the VRI point, from there it has two branches to the two next *SP*_1_, and one branch normally continues uphill to an *SP*_2_, a saddle of index two. The bifurcation explains the observed ‘hyperbolicity’ of NTs nearby a VRI in (1), an old known property.

BB claim that the green lines, the curves with Det(H)=0, are convex seen from the minima. This is correct in many cases, however, not throughout. There are staight lines in Figure 2 of reference (10), as well as a concave line in Figure 6 of (2).

## Discussion

We propose to lead back Eq.[2] to the desingularized form, the Branin equation [3] and use it consistently. Note that for Eq.[7] of (1) the essential *Det*(*H*)-factor is used, but not in the text before. Why? We propose to use [3] because we think that paper (1) examines an important topic in mechano-bio-chemistry, namely the switching behavior of reaction pathways under external force. It is an important point.

There may be a further effect. Eq.[3] is ‘robust’ against numerical deviations. A small step besides the correct NT causes another NT to a slightly other direction. For Eq.[2] we obtain after a deviation another character of the curve which now does not connect stationary points, like the magenta squiggle in the right panel of Figure 1 demonstrates.

## Conclusion

We state that the theory of NTs already offers many tools for the investigating of reaction path models. Because the solutions of [3] can just serve by itself for reaction pathway models, apart from the well known steepest descent model or gradient extremals. So also in the quite difficult field of catch bonds, we propose to use the correct NTs. A consequent application of the theory of NTs would make paper (1) easier to understand. We demand the use of the Branin equation [3] for the treatment of mechanochemical problems. We reject the claims of so called switch-points and switch-lines in (1), see the contradiction in (14).

## ACKNOWLEDGMENTS

The authors thank the financial support from the Spanish Ministerio de Economía y Competitividad, Project Nos. PID2019-109518GB-I00; Spanish Structures of Excellence María de Maeztu program, through Grant No. CEX2021-001202-M. Agència de Gestió d’Ajuts Univeristaris i de Recerca of Generalitat de Catalunya, Project No. Projecte 2021 SGR 00354.

